# Does ensembling improve feature attributions from sequence-to-activity models?

**DOI:** 10.64898/2026.07.08.737315

**Authors:** Alexandra Maslova, Maxwell W Libbrecht

## Abstract

Sequence-to-activity models take as input DNA sequence and predict genomic activities such as transcription factor binding and gene expression. Applying explainable AI (xAI) methods such as DeepLIFT to these models has recently led to breakthroughs towards many genomic problems, including transcription factor binding grammar and predicting effects of genetic variants. However, there remains significant uncertainty about the reliability of sequence-to-activity interpretations. Thus, we need accurate probabilistic measures of confidence to distinguish reliable from unreliable interpretations. Towards this end, researchers have recently aimed to characterize variability across ensembles of S2A models. However, previous work has focused on using model ensembles to improve the model predictions themselves. Here, we aim to evaluate whether model ensembles can also be used to improve feature attributions from post-hoc xAI methods. We find that ensembling attributions from multiple models improves downstream applications, including identifying transcription factor motifs and predicting regulatory genetic variants. We show that forming an ensemble using Monte Carlo Dropout (MCDropout) gets near to, but does not match, the performance of training multiple models, at much less train-time computational cost.

## 1 Introduction

A major goal of molecular biology is to decipher the code governing genomic cis-regulatory elements. Deep neural network sequence-to-activity (S2A) models are poised to help accomplish this goal. These models take as input DNA sequence and predict aspects of the regulatory activity of the sequence, such as chromatin accessibility, transcription factor binding and gene expression [32, 1, 2]. These models enable researchers to discover functional DNA elements and their biological mechanisms of action. For example, they allow researchers to quantify the impact of sequence variants on biological activity [25], and to discover the combinatorial rules and mechanisms of transcription factor binding which allow the encoding of complex cellular functions [8].

Because researchers are often interested in deriving scientific insights rather than raw predictions, they commonly use explainable AI (xAI) methods to understand the patterns a trained model has learned. Post-hoc feature attribution methods such as DeepLIFT [28] or Enhanced Integrated Gradients [15] are used to identify biologically important sequence features [20].

However, there remains significant uncertainty over the reliability of sequence-to-activity predictions [26, 13, 16] and xAI interpretations of those models [19, 29, 4]. Recent work questions the generalizability of genomics sequence-to-activity models, as demonstrated by their poor ability to predict the effects of individual genetic variation and cell-type specific activity [26, 13, 16]. Additionally, researchers showed that predictive performance is not necessarily indicative of the quality of feature attributions [18].

Given these challenges, it is important to evaluate the uncertainty in model predictions and interpretations. One way to characterize a model’s uncertainty is to collect output from an ensemble of learned models. Observing that different high-quality models disagree about predictions for a particular sequence indicates that these predictions are unreliable. These observations can be used to discount unreliable outputs, and can indicate which sequences are most valuable to experimentally assay through an “active learning” approach [12, 27, 31, 10, 24]. (Specifically, this approach evaluates *epistemic*, or model-based, uncertainty; there remains an additional layer of irreducible *aleatoric*, or data-based, uncertainty.)

For this reason, a number of researchers have recently aimed to characterize the variability of outputs from multiple S2A models. Bajwa et al. [3] trained five Basenji2 replicates with identical hyperparameters but different random seeds and showed that while reference genome predictions are highly consistent, predictions on variant sequences are unreliable. Zhou et al. [32] introduced DEGU (Distilling Ensembles for Genomic Uncertainty-aware models), which takes as input an ensemble of trained models and distills them into a single model that captures both the average and variability of the ensemble’s predictions. Researchers have also explored Monte Carlo Dropout (MCDropout) as an approach to generate an ensemble of predictions (Methods) [11, 30, 9, 24].

This suggests a simple strategy for improving interpretation reliability: aggregate feature attribution scores across multiple independently trained models. However, due to the high computational cost of training deep models, this approach may not always be feasible in practice. Here, we propose a computationally cheaper approach based on the recently-proposed method of Monte Carlo Dropout (MCDropout) which has been shown to improve the quality of model predictions as well as provide a Bayesian approximation of model uncertainty [11, 30, 9]. Rather than requiring multiple independently trained models, MCDropout approximates an ensemble by sampling model replicates from a single trained model at inference time by randomly pruning a subset of model nodes.

However, previous work has focused on using model ensembles to improve the model predictions themselves. Here, we aim to evaluate whether model ensembles can also be used to improve feature attributions from post-hoc xAI methods. We build upon the work of Bajwa et al. [3], focusing on five Basenji2 replicates. We test two complementary strategies for generating model ensembles: repeated re-training of models and Monte Carlo Dropout (MCDropout) applied at test time. In both cases, we assess whether combining attribution information across multiple model realizations improves the identification of biologically meaningful features compared to relying on a single model alone.

## 2 Results

### 2.1 Evaluating attribution quality by TF motif match

First, we aimed to characterize the degree to which attributions for predictions of transcription factor (TF) binding match that TF’s known motif. To do this, we focused on Basenji [17], a deep convolutional neural network for regulatory genomics that transforms 131,072 bp DNA sequences into predictions of thousands of experimental signal tracks summarizing transcription factor binding, chromatin state, and transcriptional activity. For the model ensemble we utilized five independently re-trained Basenji replicates produced by Bajwa et al.[3].

Because no true ground-truth attribution maps exist for the genomic data on which Basenji was trained, we used experimentally observed transcription factor binding sites and their corresponding known sequence motifs as a proxy for attribution quality. Specifically, we focused on the 1582 Basenji output tracks corresponding to ChIP-seq experiments that profile transcription factor binding. We evaluated whether high attribution scores were concentrated at positions matching the expected binding motifs within experimentally detected peak regions. To ensure sufficient signal for robust evaluation, we excluded ChIP-seq datasets with fewer than 500 peaks, leaving 201 tracks corresponding to 50 unique transcription factors.

We devised a “motif match score” quantifying the degree to which each base pair contributes to a high-confidence motif match. For a signal substantially determined by sequence motifs, we expect the signal’s attribution to be similar to the motif match score. For each track, we quantified attribution quality by measuring the agreement between base-pair-resolution attribution scores and motif match scores within 128 bp windows overlapping ChIP-seq peaks. Motif match scores were computed for all human transcription factor motifs in the JASPAR database [21] that matched the name of a TF in our dataset (Methods). We found that averaging the attributions across five models resulted in better similarity between attributions and the motif match score, suggesting that attributions better capture the motif’s effect (Figure 1). We then quantified this relationship by computing the area under the ROC curve (AUC) between attribution scores and motif match scores across all evaluated tracks. For each ChIP-seq dataset, we identified the best-matching JASPAR motif and compared the resulting AUC values for single-model and ensembled attributions.

**Figure 1:**
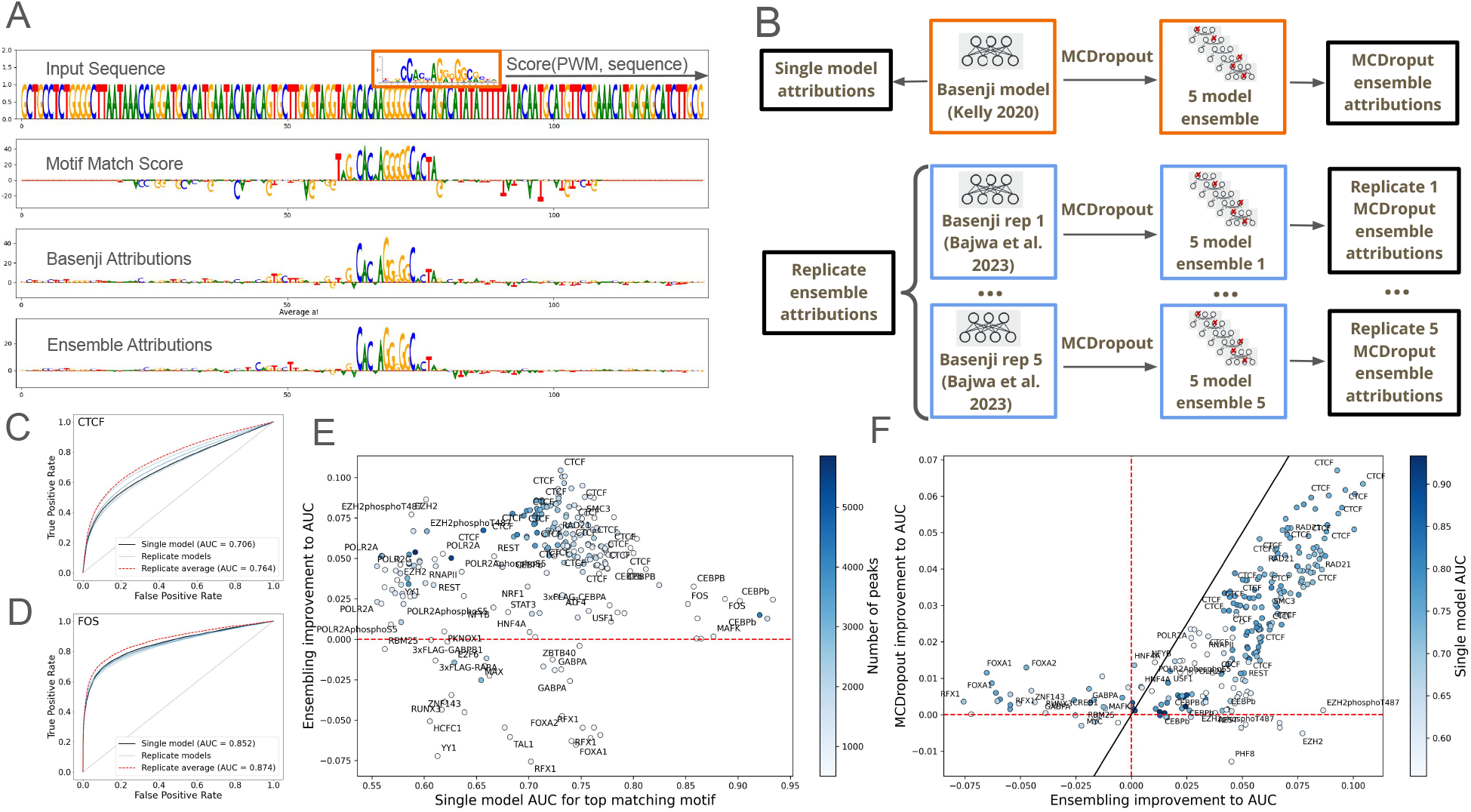
**(a)** Example of motif-resolved attribution signal at a CTCF-bound locus. First panel - position weight matrix (PWM) of the CTCF transcription factor binding motif from the ENCODE database [7]. Second panel - base-pair-level motif match scores for the CTCF motif from JASPAR[21] across the input sequence window. Panel 3 - attribution scores from a single Basenji model for the same locus. Panel 4 - attribution scores obtained by averaging base-pair-level attributions across five independently re-trained Basenji replicates. **(b)** Overview of models used in the Basenji analysis, including all ensembles. **(c-d)** ROC curves evaluating the degree to which model attributions (with or without ensembling) captures the motif match score for representative CTCF and FOS ChIP-seq tracks. **(e-f)** Comparison of AUC improvements from ensemble attribution methods. **(e)** The improvement to AUC when using averaged attribution values from 5 different basenji models versus a single basenji model, as a function of the original AUC value for the best-matching motif. The hue corresponds to the number of unique binding events identified in each dataset and used for the AUC calculation. **(f)** The improvement to AUC (calculated as difference in AUC values) of attribution values averaged from a retrained model ensemble (x-axis) versus MCDropout model ensemble (y-axis). All results are computed for best matching motif in the JASPAR human motif database, as determined from AUC values for the base Basenji model. Plot shows all human ChIP-seq output predictions from that Basenji human output head that have at least 500 identified peaks within the test dataset.

Ensembling improved attribution AUC for 175 (or 87%) of the 201 tested ChIP-seq datasets, indicating that aggregating attributions across independently trained models yields more biologically meaningful interpretation maps than relying on a single model alone.

We next asked whether the improvements observed in re-trained model ensembles could be replicated with a computationally cheaper MCDropout model ensemble. To test this, we applied MCDropout to the original Basenji model [17] as described in Methods to that obtained from the ensemble of independently retrained Basenji models.

Overall, MCDropout improved attribution quality relative to the single-model baseline and broadly recapitulated the gains observed with retraining-based ensembles (Figure 1f). Tracks with larger improvements from retraining also tended to show larger improvements from dropout-based averaging, indicating that MCDropout recovers much of the same stabilizing effect on attribution scores. Although retraining remained somewhat stronger for some tracks, these results suggest that MCDropout provides a useful low-cost alternative for improving attribution quality in Basenji.

#### Ensembling improves attribution quality in simulated motif-grammar tasks

To confirm these findings in a setting where the true informative sequence features are known, we simulated a heterodimer binding task (References methods). In these simulated datasets, class labels depend on the presence of two motifs in a specific spatial arrangement, allowing attribution performance to be evaluated directly against the ground-truth embedded motif positions (References methods).

For each of the two simulated heterodimer tasks, we trained 10 independently initialized model replicates and then generated 10 MCDropout realizations from each retrained model, producing a total of 100 dropout-based model realizations per task. Attribution quality was quantified by comparing base-pair attribution scores to motif match scores at the embedded motif positions using ROC AUC. Because the simulated labels are generated from known motif configurations, this framework provides a direct benchmark of whether attribution ensembling improves the recovery of the true regulatory grammar.

For both simulated data experiments, averaging attribution scores across retrained models improved attribution accuracy relative to individual model replicates, and averaging across the larger MCDropout ensemble improved performance further (Figure 2c-d). These results indicate that stochastic averaging suppresses model-specific attribution noise and yields attribution maps that more closely recover the known motif instances responsible for model output.

**Figure 2:**
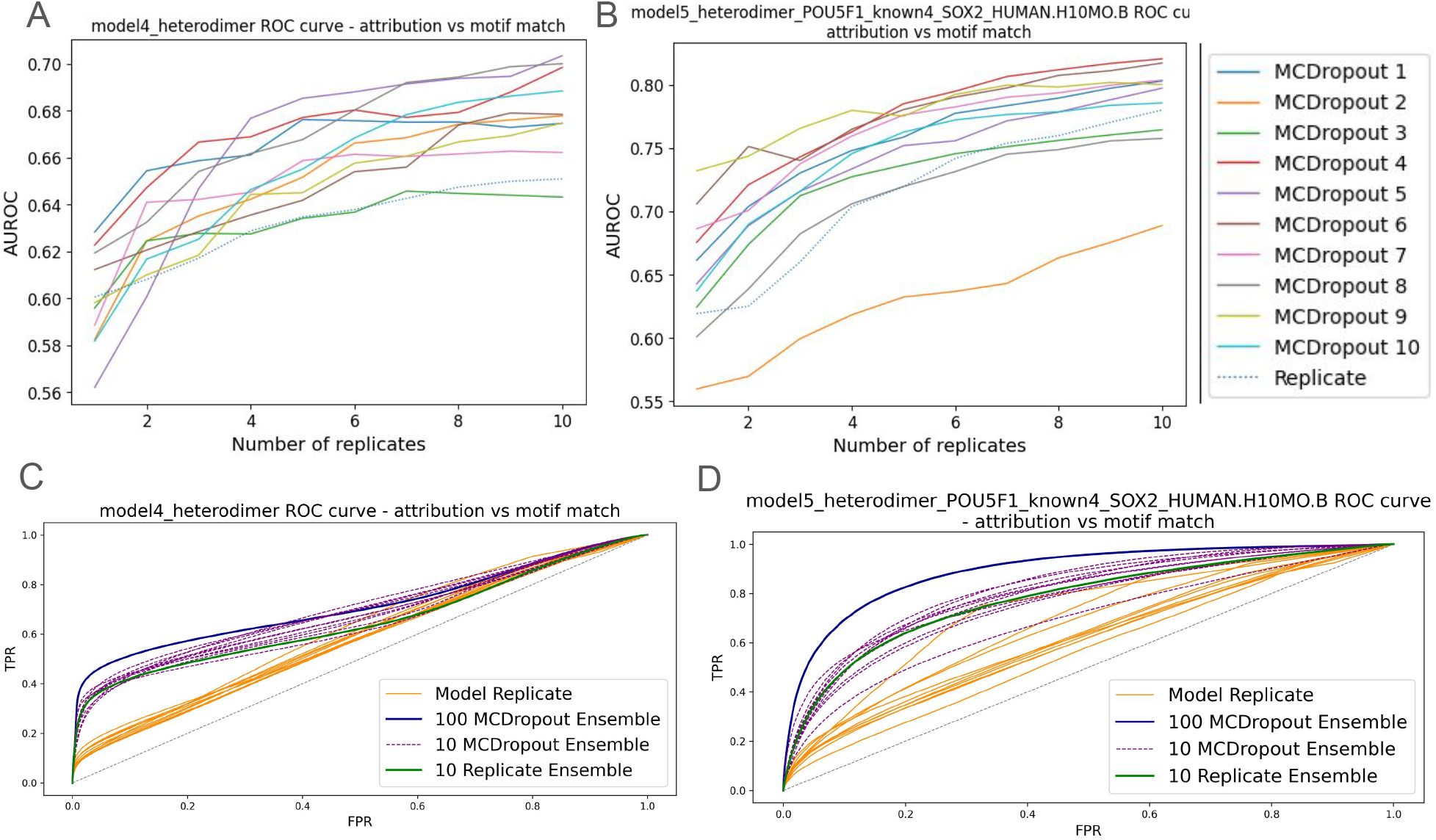
Accuracy of attribution values derived from models trained on simulated data under two heterodimer scenarios. The performance is measured by the concordance between attribution values and motif match values for the motifs embedded in the simulated data at base-pair resolution. **(a-b)** AUC for ensembled model attribution values as a function of the number of models in the ensemble. The experiment was performed with the 10 re-trained model replacates and for all 10 sets of MCDropout models create from each of the 10 re-trained models. **(c-d)** For each model, the plots show the performance of 10 independently retrained replicate models, the ensemble average across retrained models, and the ensemble average across 100 MCDropout models generated from the retrained replicates.

We also evaluated how attribution performance depends on ensemble size. In both simulation settings, attribution AUC increased as more models were included in the ensemble, with the MCDropout ensembles providing a practical way to scale ensemble size beyond what would be feasible with retraining (Figure 2a-b). These results also indicate that the benefit of additional MCDropout models becomes marginal as more are added, suggesting diminishing returns beyond moderate ensemble sizes and indicating that much of the gain in attribution accuracy can be captured with a relatively small number of stochastic samples.

### 2.2 Evaluating ensemble performance for predicting effects of quantitative trait loci

To evaluate whether MCDropout-based attribution improvements translate to practical benefits for genomic variant interpretation, we assessed attribution quality on quantitative trait loci (QTL) benchmark prediction tasks established in DART-Eval [23]. This benchmark consists of two chromatin accessibility QTL datasets: “African caQTLs” derived from ATAC-seq measurements of lymphoblastoid cell lines (LCLs) from six different African populations, and “Yoruban dsQTLs” derived from DNase-seq measurements of 70 Yoruba-derived LCLs. These datasets provide real-world benchmarks where we can directly assess whether ensembling correlates with better predictions and attributions of genetic variant effects.

We selected chromatin accessibility tracks from the Basenji training set that matched the cell type backgrounds of each QTL study as described in Methods, resulting in 21 EBV-transformed lymphocyte tracks that most closely align with the LCLs used in the African caQTL and dsQTL experiments. For each QTL dataset, we compared the ability of single-model, as well as retrained replicate and MCDropout ensembles to accurately predict the experimentally observed change in chromatin accessibility for the reference versus alternative alleles. Additionally, we evaluated how well the attribution values at the reference allele of each QTL correlate with the QTL effect size.

Ensembling retrained replicates improved the correspondence between predicted and experimentally observed QTL effects, and likewise improved the agreement between the attribution assigned to each SNP and its observed effect size (3b). This benefit was specific to retrained replicate ensembles: MCDropout ensembles did not reproduce it, yielding no gain in either predictive accuracy or attribution correspondence relative to a single model.

We next asked whether the variance in attribution values across replicates could serve as a per-SNP indicator of attribution reliability. We reasoned that if inter-replicate variance reflects genuine model uncertainty, then SNPs whose attributions vary most across replicates should also be those whose attributions correspond most poorly to the observed QTL effect. To test this, we fit a linear model relating the reference-allele attribution to the observed QTL effect for each dataset (Figure 3c-d) and examined the relationship between each SNP’s inter-replicate attribution variance and its residual from that fit. Attribution variance was positively associated with residual magnitude across all five MCDropout ensembles and the 5 re-trained replicate ensemble (Figure 3e), indicating that both are informative for capturing model attribution uncertainty.

**Figure 3:**
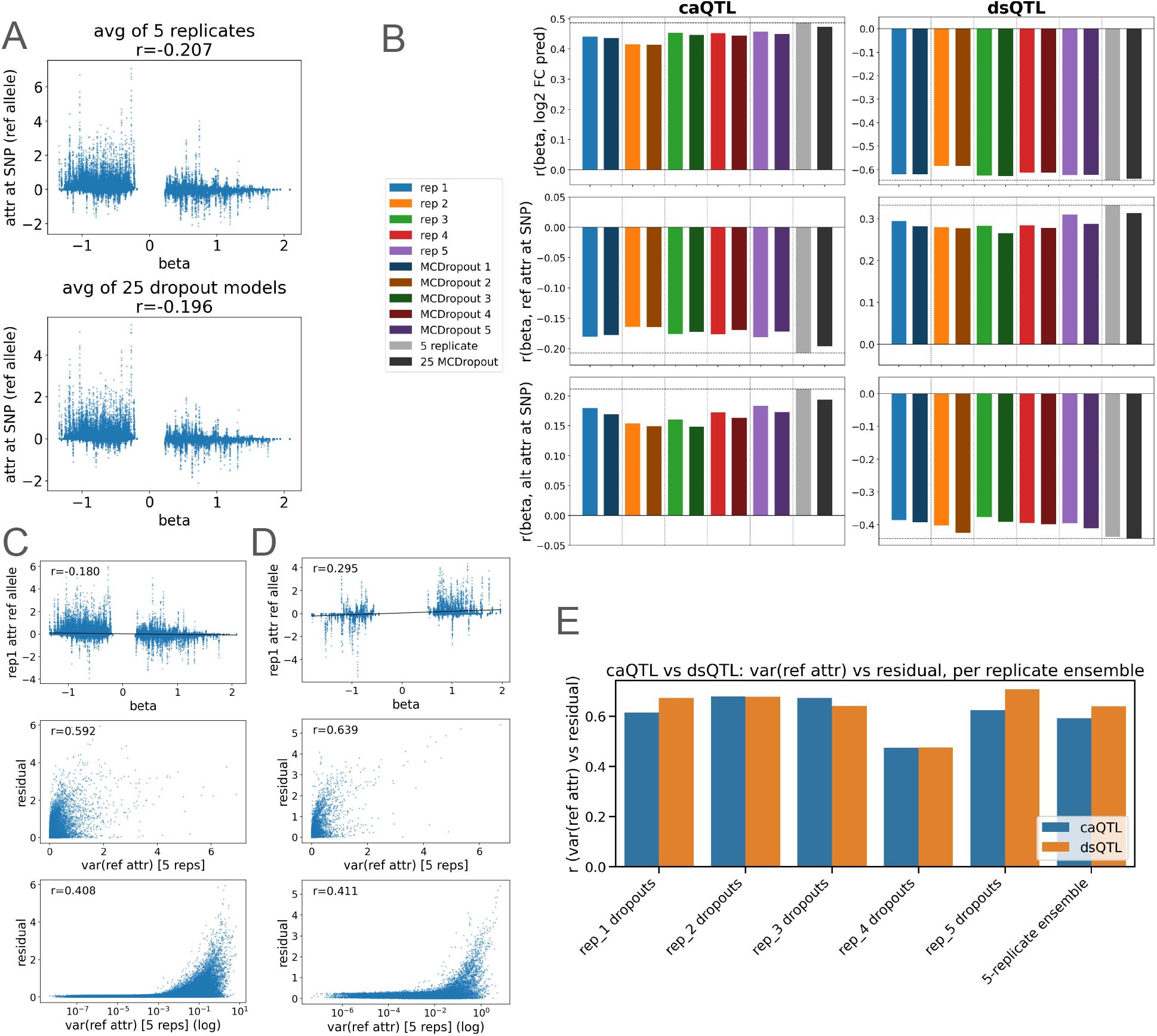
**(a)** Degree to which ensembled attributions (vertical axis) capture chromatin accessibility QTL (beta, horizontal axis). **(b)** Pearson correlation between the predicted log fold change difference in predictions, attribution at reference and attribution at alternative allele for all 5 model replicates and MCDropout ensembles. The reported QTL beta statistic of the caQTL and dsQTL data sets have opposite sign relative to one another, leading to opposite signs in all analyses. **(c-d)** First panel shows a linear fit obtained for each attribution versus QTL effect plot. Second panel shows the variance in SNP attribution values across 5 replicate models versus the linear fit residual. The third panel shows the variance values on a log scale. **(e)** Correlation between reference allele attributions and linear fit residuals across all 5 MCDropout ensembles and the retrained replicate ensemble.

### 2.3 MCDropout recapitulates the attribution uncertainty of retrained ensembles

We next asked whether the uncertainty captured by MCDropout ensembles reflects the same underlying model uncertainty as that captured by independently retrained ensembles. If so, MCDropout could serve as a computationally inexpensive way to derive a measure of uncertainty.

We evaluated this concordance in two complementary settings. In the first, we examined ensemble variance for all six MCDropout ensembles, computing the variance of reference-allele attributions across all QTLs and their relevant output tracks (Figure 4a-b). In the second, we examined attribution variance in the TF binding dataset. Using the same 201 ChIP-seq binding tracks described previously, we computed the attribution variance at every base pair across all 128 bp sequences containing a binding event, yielding a dataset that spans both high-importance positions (those overlapping the binding motif) and low-importance positions (Figure 4c).

**Figure 4:**
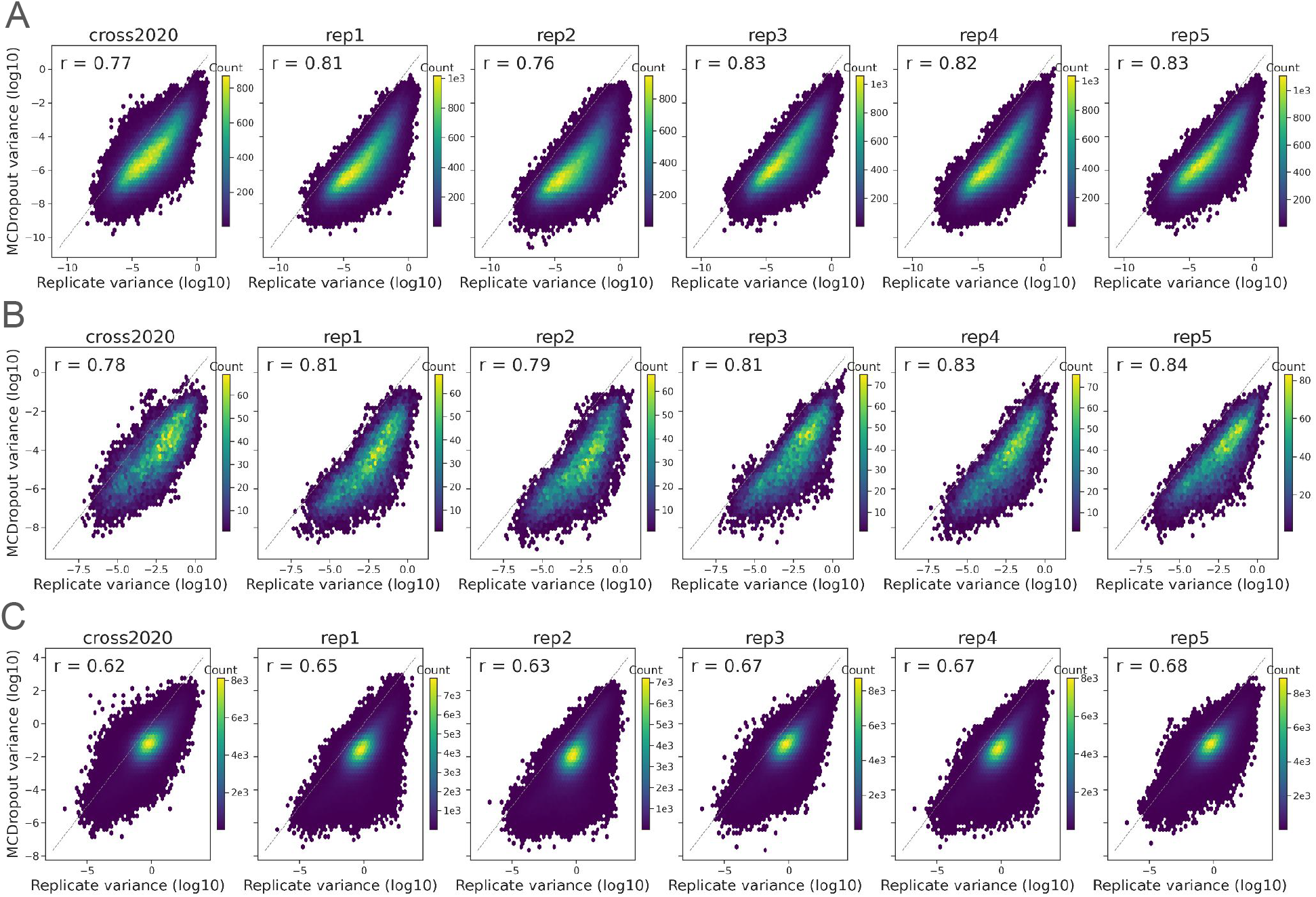
Variance across model ensemble attribution values for **(a-b)** each QTL reference allele in 21 matching cell types and (**c**) all positions within 128bp bins containing ChIP-seq peaks across all TF binding datasets in the Basenji test set containing more than 500 peaks total.

Across both settings, the per-position attribution variance from the MCDropout ensembles was positively correlated with the variance from the retrained replicate ensemble, indicating that MCDropout is able to recover much of the same model uncertainty as retraining. Notably, however, the retrained ensembles exhibited consistently higher attribution variance than the MCDropout ensembles. This is expected, as retraining perturbs the full set of learned weights and therefore introduces greater variability across model realizations than stochastically masking activations within a single trained model.

## 3 Conclusions

Here, we characterized the utility of model ensembles for S2A feature attributions. We showed that averages across ensembles of models generally improve downstream applications of feature attributions, including identifying TF binding sites and QTLs. Because MCDropout can be applied to a pre-trained model, it is usable for low-resource groups applying models trained by high-resource groups who did not train (or did not share) multiple trained models. We found the MCDropout’s performance approaches, but does not match, that of independent models.

There remains significant future work in this area. We focused here on Basenji, as it is a large, high-performing model. It remains to be shown how these results extend either to small models or the most recent ultra-large models.

Because the number of parameters of a model influences the bias-variance tradeoff, differently sized models may behave significantly different in reponse to ensembling.

## 4 Methods

### 4.1 Sequence-to-activity models and epigenomics data

#### Simulated experiments

We performed the simulated data experiments utilizing the datasets and model architectures described in [14].

Each dataset contains 20,000 sequences of length 500 base pairs, split evenly between positive and negative classes. We generated synthetic sequences using the simdna Python package (https://github.com/kundajelab/simdna). The background sequence is sampled uniformly at random, while motifs are embedded according to position weight matrix (PWM) probabilities. When embedded, motifs are inserted in either the forward or reverse-complement orientation with equal probability (0.5), reflecting strand symmetry in real regulatory DNA.

For our experiments, we generated two binary heterodimer motif datasets designed to capture simple motif grammar. Each sequence contains two distinct transcription factor motifs embedded at random positions. Negative sequences contain the two motifs placed independently at arbitrary locations. Positive sequences contain the same two motifs embedded in close proximity, with an inter-motif spacing of 2–5 base pairs. The task requires the model to distinguish isolated motif occurrences from a specific spatial configuration indicative of cooperative binding. We tested the approach with two sets of motifs: SPI1, IRF and POU5F, FOX2.

We used the following simple model architectures for this predictive task:

|  |  |
| --- | --- |
| **SPI1, IRF**<br>- 15 conv, size 10, stride 1, dropout 0.2<br>- 15 conv, size 10, stride 1, dropout 0.2<br>- 15 conv, size 10, stride 1, dropout 0.2<br>- max-pooling size 35, stride 35<br>- fully connected output layer | **POU5F, FOX2**<br>- 30 conv, size 10, stride 1, dropout 0.2<br>- 15 conv, size 10, stride 1, dropout 0.2<br>- 15 conv, size 10, stride 1, dropout 0.2<br>- 15 conv, size 10, stride 1, dropout 0.2<br>- max-pooling size 35, stride 35<br>- fully connected output layer |

To train the models, we used a randomly selected 80 percent of the examples as a training and validation set and the rest as the test set. We trained both models for 200 epochs using the Adam optimizer with a learning rate of 0.001 and binary cross-entropy loss. We computed the validation set loss at the end of each epoch and saved the weights yielding the lowest validation loss as the final model.

##### 4.1.1 Basenji model experiments

For the genomic sequence-to-activity experiments, we used the human prediction head of Basenji2 as the primary model [17]. Basenji2 is a large multi-task convolutional neural network that takes 131,072 bp DNA sequences as input and predicts regulatory activity across thousands of genomic tracks at 128 bp resolution.

All Basenji experiments were performed on the original Basenji2 human test set, corresponding to the same data split used for the retrained models. This test set consists of 1,936 genomic sequences, each 131,072 bp (2^17^ bp) in length, sampled from the human genome as described in Kelley et al., 2020 [17]. To evaluate attribution quality, we focused on transcription factor binding ChIP-seq tracks present in the Basenji training targets. Specifically, we considered the 1,583 output tracks corresponding to TF binding ChIP-seq experiments.

For each ChIP-seq experiment, we identified all 128 bp output bins in the test set containing a positive TF binding event, defined as a target signal greater than 60 in the corresponding ChIP-seq track. Because robust attribution evaluation requires a sufficient number of positive sites, we excluded tracks with fewer than 500 detected peaks in the test set. After filtering, 249 ChIP-seq TF binding tracks remained for downstream analysis.

### 4.2 Feature attribution

Feature attributions in deep neural networks refer to the techniques used to interpret and understand the contributions of input features to the network’s predictions. In the context of genomics, feature attributions can help identify which specific parts of a DNA sequence are most influential in predicting genomic activity. These sequence features are then used to form hypotheses about which parts of the DNA sequence encode the biological programming of the interrogated cell types. Our approach focuses on these gradient-based methods for obtaining feature attributions because they be efficiently computed through a single back-propagation pass through the network, meaning we can apply them to multiple test samples and models without too many computational resources.

For the simulated data the per-position attribution scores were obtained across the test sequences using DeepLIFT (Deep Learning Important FeaTures)[28]. We utilized the captum package for Python (available at https://github.com/pytorch/captum) to compute these attribution scores. Because the attribution value signal was dominated by a small number of outlier values, we clipped the attributions at the 1st and 99th percentile values for large negative and positive attributions, respectively.

For the Basenji model experiments, nucleotide-resolution attribution scores were computed using gradient *×* input, following [17]. Specifically, given an input sequence represented as a one-hot encoded matrix, attribution at each base was defined as the element-wise product between the input and the gradient of the selected output with respect to that input. We used this method in place of DeepLIFT because the large receptive field and model size of Basenji make DeepLIFT computationally difficult, and because it has been used for Basenji in previous work. The resulting attribution values provide a position-specific estimate of how strongly the observed nucleotide contributes to the output prediction for the target assay.

### 4.2 Ensembling models and MCDropout

To evaluate ensemble-based attribution strategies, we considered two types of model variation. First, we used five independently retrained Basenji models from Bajwa et al. [3]. Second, we generated five MCDropout variants from the original Basenji2 model as well as for each of teh 5 re-trained replicates, resulting in a total of 6 MCDropout model ensembles, as depicted in Figure 1b. We did so by enabling dropout at inference time, using the same dropout rate specified for each layer during training. A drop-out rate of 0.3 was applied to all eleven convolutional layers in the dilated residual block, and a drop-out rate of 0.05 was applied to the convolutional layer preceding the final linear layer.

### 4.4 Evaluations

#### 4.4.1 Motif scanning

We posit that an effective attribution method for capturing transcription factor binding should assign high importance to base pairs corresponding to a transcription factor motif within sequences corresponding to positive binding events. We therefore devised a method to measure a per-position motif similarity score using a motif PWM as input that can be used to evaluate the quality of attribution values at each base pair.

Each base pair within a test sequence is assigned a score based on the best alignment of the motif at that position and the information content of the aligned position within the position weight matrix(PWM). First, for some position *i* within the test sequence *S* the best alignment of the target PWM is found by maximizing the value of the convolution between the PWM and sub-sequence of *S*. Then, given this alignment, an individual base-pair score is assigned based on the weight of the particular base at the aligned position within the PWM.

Formally, for position *i* within a 4*xL* matrix representing a one-hot-encoded DNA sequence *S*, and a position weight matrix *P* of length *l*:

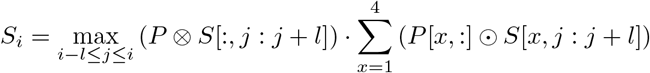

For the Basenji model experiments, motif scanning was performed using human transcription factor motifs from the JASPAR database [21].

#### 4.4.2 Chromatin accessibility QTL datasets

To further evaluate feature attribution values in a biologically-relevant context we used two chromatin accessibility quantitative trait loci(QTL) benchmark datasets presented in DART-Eval [23].

The first dataset consists of chromatin accessibility QTLs (caQTLs) reported by de Gorter et al. [6], who profiled accessibility by ATAC-seq in lymphoblastoid cell lines drawn from six African populations represented in the 1000 Genomes and HapMap projects. The full set comprises 219,832 variants annotated with effect sizes and significance values. We further filtered the data by restricting the set to significant SNPs lying within *±*100 bp of the top 50,000 peaks across all 100 individuals as well as possessing pre-computed Enformer scores, as described in Pamapri et al.[22], yielding a total of 6,821 caQTLs for this dataset.

The second dataset consists of DNase I sensitivity QTLs (dsQTLs) reported by Degner et al. [5], who measured chromatin accessibility by DNase-seq in 70 Yoruba lymphoblastoid cell lines with available genome-wide genotypes and mapped dsQTLs by associating accessibility within each DNase I hypersensitive window with nearby genotypes across a surrounding 40 kb region. Following the processing of Pampari et al. [22] described above for the caQTL dataset, we retained 560 significant dsQTLs for analysis.

To identify Basenji output tracks relevant to these LCL-derived variants, we matched the cell types and biosamples used in the Basenji training data to GTEx v11 tissue annotations downloaded from the Genotype-Tissue Expression (GTEx) Project web portal (https://gtexportal.org/). We performed this matching with the assistance of Claude (Anthropic) using the following prompt: “Files ENCODE_celltypes.tsv and GTEX_celltypes.csv each contain a list of cell types, tissues and biosamples used by ENCODE and GTEX respectively. For each row in ENCODE_celltypes.tsv, please choose the GTEX cell type that best matches it. Output this to a file matched_celltypes.csv containing (1) the ENCODE cell type, sthe matched GTEX cell type, and (3) a brief summary of the reasoning you used to make this match. Use your best judgement, aiming to match the cell types which are most similar biochemically.” From the resulting annotations, we selected the 21 accessibility tracks labeled as EBV-transformed lymphocytes, which provided the closest tissue match to the LCLs used in both QTL experiments.

